# Stably coexisting communities deliver higher ecosystem multifunctionality

**DOI:** 10.1101/2025.11.21.689673

**Authors:** Caroline Daniel, Oscar Godoy, Noémie A. Pichon, Mathis Gheno, Eric Allan

## Abstract

Linking the processes determining species coexistence and effects of biodiversity on ecosystem functioning is important for a mechanistic understanding of how species loss impacts ecosystems^1^. There are good theoretical reasons to expect stably coexisting communities to deliver higher ecosystem functioning, as niche differences should promote coexistence and complementarity^2^, however, there is still little empirical evidence to demonstrate this relationship^3^. We present a novel experimental approach to tackle this question, in which we first measured interaction coefficients and intrinsic growth rates for 12 grassland species^4^ and used population models to estimate coexistence^5^. We then assembled 48 three-species communities, predicted to differ in their degree of coexistence and measured seven above and belowground functions. Here we extend findings demonstrating that biodiversity increases multifunctionality^6,7^ to show that amongst communities with the same initial diversity, multifunctionality is higher when all the species are predicted to coexist. Decomposing coexistence into the influence of niche and fitness differences showed that multifunctionality was high in niche differentiated communities but also in those with a single highly competitive species. We further examined the effect of indirect interactions, which provide an alternative mechanism of coexistence in multispecies communities, and found that they also provided an alternative route to high multifunctionality. Our results point the way to a unification of coexistence and biodiversity theory and to a more mechanistic understanding of the functional effects of biodiversity change.

## Main

Decades of experimental research has overwhelmingly shown a positive causal relationship between biodiversity and ecosystem functioning^8,9^, with diverse communities providing a larger number of functions at a high level^10,11^. Biodiversity may increase levels of particular functions through complementary effects, which can arise, for instance, when species differ in resource use or when they protect each other against herbivores and pathogens^12^. The same underlying processes can drive stable niche differentiation, and therefore communities assembled from species that can stably coexist should have higher functioning than communities in which not all species are expected to persist in the long-term^1^. However, very few studies have connected the mechanisms maintaining species diversity with effects on ecosystem functioning^3,13^ and none have linked coexistence mechanisms with the ability of a community to simultaneously supply multiple functions. For diversity to drive high levels of multifunctionality, we expect multiple functions to increase with diversity and/or for species to complement each other by supplying different functions^6,7,10^. On the other hand, if one or a few species can provide a high level of functioning on their own, increasing species richness would not increase their multifunctionality. If we can identify a link between the ability of a community to coexist and its ability to deliver high levels of multifunctionality, we would be able to better understand the mechanisms underlying the indirect impact of global change on ecosystem functioning^14^.

To unravel the links between coexistence and multifunctionality, we need to connect recent, theoretical measures of the extent to which multispecies assemblages can coexist, with novel experiments. The degree of multispecies coexistence can now be quantified as a continuous variable, rather than a dichotomous outcome, by calculating how much perturbation a community can undergo before one species goes extinct^15^. Theory predicts that communities with “strong” species coexistence can withstand large perturbations without losing species because there is an “excess of niche differences” compared to the minimum required to coexist, i.e., the minimum niche differences required to overcome the fitness differences (also called competitive ability differences)^16,17^. Empirical evidence shows that species pairs with stronger coexistence promote higher levels of biomass production^3^, however, it is unclear whether this result extends to multiple functions. We might expect that larger niche differences would promote higher multifunctionality because species that differ in their environmental response would also supply different ecosystem functions, e.g., species differing in their competitive abilities for nutrients or light could differ in productivity and in effects on nutrient cycling^18^. However, high niche differences might reduce other functions, like energy flow to higher trophic levels, if they result in reduced sharing of herbivores or pathogens between plants^19^. In contrast, fitness differences, measured as disparities in species’ intrinsic growth rates, do not promote coexistence but rather the dominance of superior competitors^20^. Nevertheless fitness differences could enhance multifunctionality if species with the highest growth rates support the most functions^21^. We could therefore expect that both high niche differences and high fitness differences would promote multifunctionality. This makes it hard to predict whether the conditions for strong coexistence, i.e., niche differences most strongly exceeding fitness differences, would lead to the highest multifunctionality. Further, it is challenging to explicitly test whether the communities that can more strongly coexist display higher levels of ecosystem multifunctionality, using classic biodiversity experiments. To solve this limitation, we need to move away from how experiments measuring multiple functions have been traditionally designed. Here, we advocate that rather than maximising variation in species richness, we need to maximize variation in the species differences that predict species coexistence (i.e., niche and fitness differences). This means creating experimental communities that vary in their predicted niche and fitness differences and in their strength of coexistence, rather than in their species number.

Attempts to empirically link coexistence and functioning have so far been limited to communities of only two species^3^. They have therefore ignored indirect effects such as indirect facilitation, indirect competition or intransitive loops that emerge from interaction chains and which could play a role in stabilising coexistence in multispecies assemblages^22^. There is little theory on how indirect interactions might affect functioning but if they allow coexistence without pairwise niche differences then indirect interactions could decouple the relationship between stable coexistence and functioning^22^. The structuralist approach^5^ is an existing theoretical framework that allows an overall quantification of the importance of these indirect interactions in affecting the strength of coexistence. Linking measures of structural niche and fitness differences, as well as indirect effects, to multifunctionality is therefore important to link coexistence and functioning in communities with more realistic levels of diversity. New measures of multifunctionality may also be necessary to explore this link. Previous studies have calculated multifunctionality as the number of functions that exceed a critical threshold level^23,24^, which allows a comparison of multifunctionality between communities. However to link multifunctionality to multispecies coexistence, we need an explicit measure of the gain in functioning for diverse communities relative to monocultures. To address this gap, we can calculate multifunctionality based on net effects of diversity on individual functions^25^ (“net effect” multifunctionality, Fig. S1). If coexistence mechanisms, such as the indirect interactions that emerge in multispecies communities, are important for increasing multifunctionality, we might expect stronger effects of indirect interactions on a measure of net-effect multifunctionality. Testing this link is critical to determine whether the theoretical conditions for multispecies diversity maintenance relate to high ecosystem functioning.

Here, we developed a new experimental approach to investigate how the mechanisms determining the strength of species coexistence affect ecosystem multifunctionality. We first conducted a competition experiment between a set of 12 perennial species (Table S1), by planting phytometer plants into different con- and heterospecific neighbourhoods, as well as into neighbourhoods with no other individuals^4^. With this experiment, we obtained information on species intrinsic growth rates and all interaction coefficients, which we coupled with a population model to predict niche differences, fitness differences, indirect interactions and hence the coexistence strength of all combinations of 3 species (triplets) from the pool of 12. We then selected 48 of these triplets, in which differences in the strength of coexistence was either due to variation in niche differences or indirect interactions. We next established 48 new experimental communities containing these triplets, together with all 12 monocultures, and grew the communities under similar climatic conditions to the first experiment, for two years (Fig. 1a). After this, we measured seven above and belowground functions and calculated multifunctionality and net effect multifunctionality (Fig. S1). To test our hypothesis that the predicted strength of coexistence in each experimental community, summarised in the minimum distance to extinction, affects multifunctionality (Fig. 1b), we fitted multimembership models to correct for shared species across communities (see methods). We further tested the relative importance of each coexistence mechanism in affecting multifunctionality (Fig. 1c), i.e., the individual effects of predicted niche differences, fitness differences, and indirect interactions. We also tested an interaction between niche and fitness differences, as we expected that high niche differences coupled with low fitness differences would increase multifunctionality (niche complementarity) but also that low niche differences and high fitness differences might promote multifunctionality, if particular species drive high levels of multiple functions. We also included an interaction between niche differences and indirect interactions to explore whether high levels of indirect interactions might reduce the positive expected relationship between niche differences and multifunctionality.

**Figure 1:**
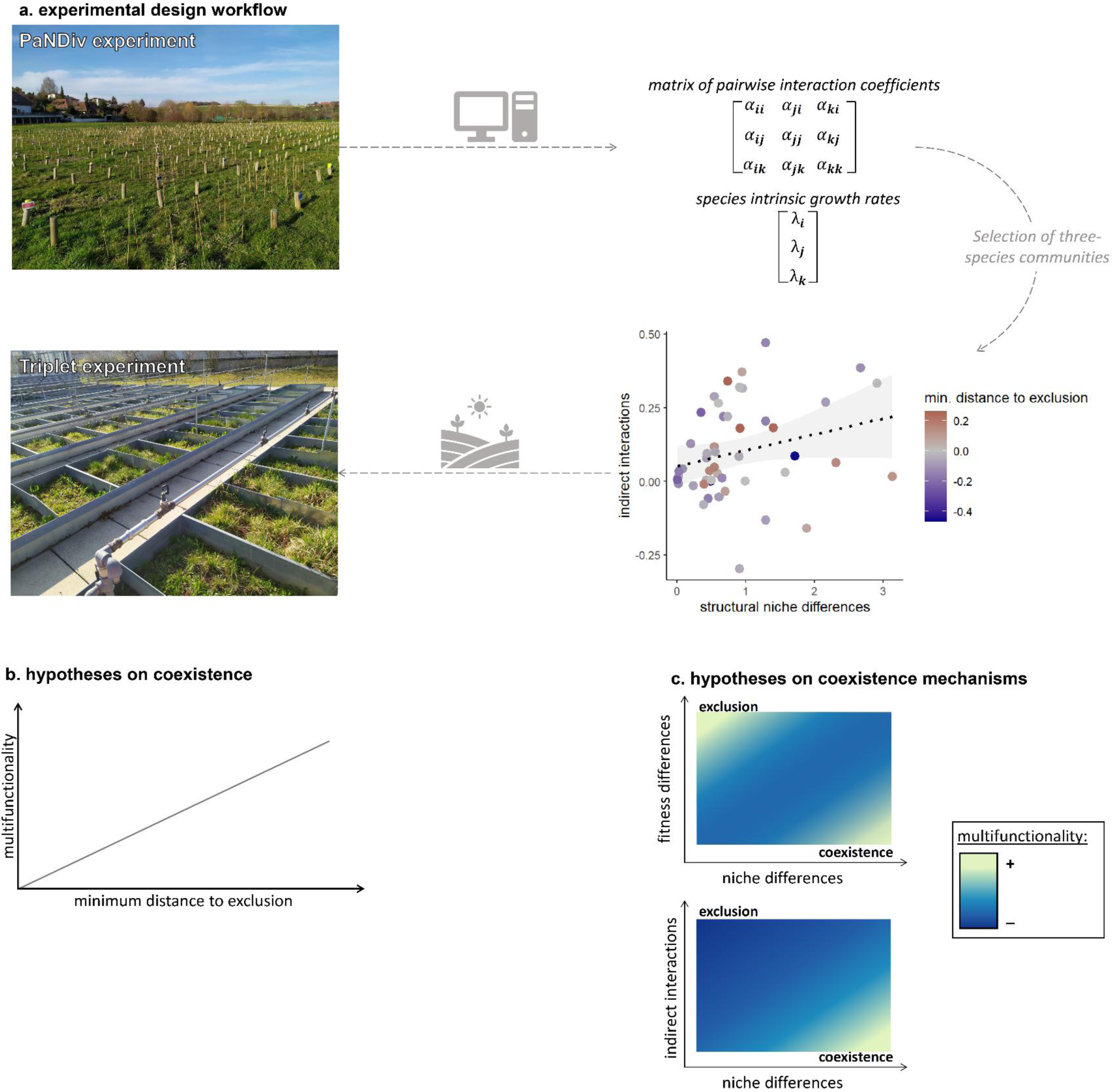
The experimental approach to test effects of coexistence on multifunctionality and the hypotheses. (a) The experimental design of this study: we first measured the response of 12 species to different neighbourhoods in the PaNDiv experiment, then computed matrices of interaction coefficients and intrinsic growth rates and selected a set of 48 triplets varying in niche differences, indirect interactions and coexistence strength. We then set up a new experiment in outdoor mesocosms, to test the effect of coexistence mechanisms on the multifunctionality of our experimental communities. Theory provides two main hypotheses (b) we expect more stably coexisting communities, with higher minimum distance to exclusion, to have higher multifunctionality, and (c) we also expect that multifunctionality should be highest in two areas: when niche differences are high but fitness differences (also called competitive ability differences) are low, and when fitness differences are high and niche differences are low. Lastly, we expect that niche differences promote multifunctionality, but only when indirect interactions are low, because indirect interactions allow coexistence without niche differences and therefore disrupt the relationship between niche differences and multifunctionality.

### Result and Discussion

In agreement with our main hypothesis, we found that multifunctionality was highest in the communities predicted to coexist most strongly (Fig. 2a and S2a). In other words, the larger the predicted minimum distance to exclusion in three species communities, the larger their observed multifunctionality. Here we used a continuous measure of coexistence strength, which links directly to predictions of population dynamics: the more positive the minimum distance to exclusion, the more positive species population growth rates are predicted to be, and the longer it should take for any species to go extinct. Multifunctionality was calculated at thresholds from 40-80%, and we estimated average effects across the thresholds, following Pichon et al. 2024^7^, to provide an integrated measure of the effect of coexistence strength. Analysing thresholds individually (Fig. S3) revealed that coexistence strength always had positive effects on multifunctionality. This highlights that an overall measure of coexistence strength, that summarises different potentially conflicting coexistence mechanisms, predicts the ability of a community to deliver many functions. Previous work has shown that higher diversity levels are required to reach higher levels of multifunctionality^6,26^, but our results demonstrate that even among communities of the same diversity level, the ones that are more strongly coexisting provide higher levels of multifunctionality. This also extends results found for aboveground biomass and species pairs^3^ to multispecies communities and multifunctionality. The fact that it was the most strongly coexisting communities that provided the highest multifunctionality suggests that a reduction in coexistence strength might reduce multifunctionality even before species go extinct^15^. Our results therefore show that it is the communities that are expected to be able to withstand larger environmental perturbation without losing species that are also able to supply many different functions.

**Figure 2:**
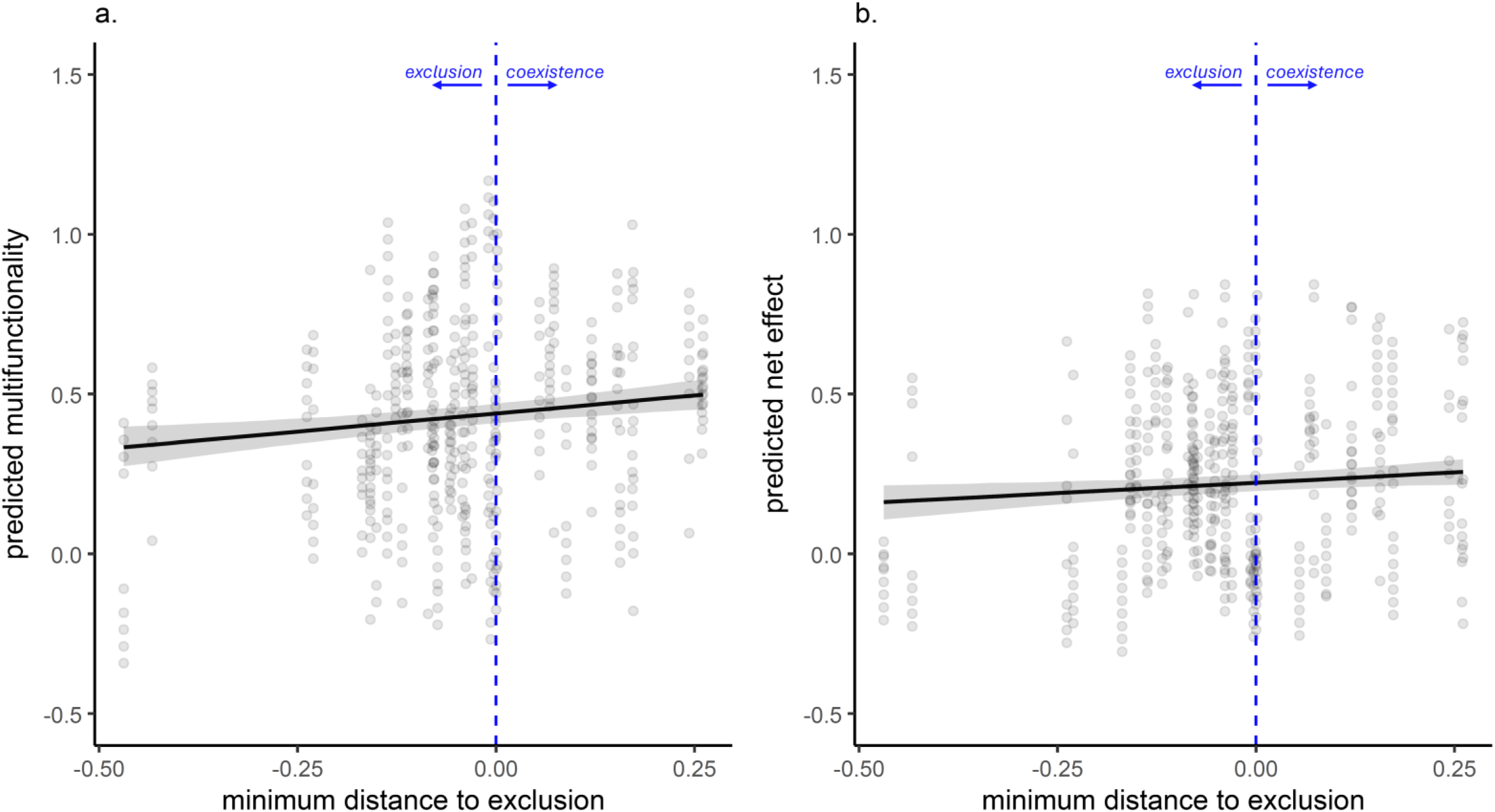
Multifunctionality (a) and net effect (b) are higher in communities predicted to coexist more stably. The black line shows predictions from multimembership models, each dots represents the residuals for each plot and each multifunctionality or net effect threshold. Minimum distance to exclusion in control conditions is derived with a structural stability approach, see methods. If the minimum distance to exclusion is positive (on the right side of the blue dotted line), the three species in the plot are predicted to stably coexist.

In line with our expectations, we also found that communities predicted to have stronger coexistence had higher multifunctionality than their monocultures (Fig. 2b and S2b), i.e., higher net effect multifunctionality (Fig. S1). On average, we found positive values of net effect multifunctionality, with 78% of functions having greater values in three species communities than predicted from their monocultures (Fig. S4). This extends the finding that diversity increases multifunctionality^10,27^ to show that amongst communities with the same number of species, diversity has the most positive effects on multifunctionality in communities where the species can most strongly coexist. This finding was possible thanks to our novel experimental approach of assembling communities differing in their predicted degree of coexistence. Our analyses therefore also show that coexistence predictions based on simple population models, and using interaction coefficients measured in a different experiment (Fig. 1), can predict levels of multifunctionality in newly assembled experimental communities. These results establish an important link between the theoretical conditions for stable coexistence and high multifunctionality. Such a link is expected from models on biomass^28–30^ and is implicit in the finding that high biodiversity promotes high functioning^18^. However, our results empirically show that the basic conditions for stable coexistence link to ecosystem multifunctionality, which points the way towards the integration of coexistence theory and biodiversity functioning research.

Despite finding significant positive effects of coexistence strength on multifunctionality, the effects on individual functions were weak. Surprisingly, we found no effect of minimum distance to exclusion on aboveground biomass production or any individual function, except for beta-glucosidase activity (Fig. S5). These findings suggest that effects of stable coexistence on multifunctionality were not driven by biomass alone and rather emerged from small effects on multiple individual functions^31^. The mechanisms that lead to overall stable coexistence may therefore not link strongly to any particular function, instead different sets of species might stably coexist through different mechanisms, similar to the finding that multivariate functional trait distances predict niche differences better than single traits^32^. However, we did not need to combine many functions, as positive effects of coexistence strength were very consistent across different sets of functions and emerged as soon as two or more functions were combined in the measure of multifunctionality (Fig. S6). The effects of coexistence strength also slightly increased the more functions were included, suggesting that strongly coexisting communities might generally supply many functions at a high level^33^. Therefore, although predicting high levels of particular functions may be challenging with overall measures of coexistence strength, it does seem that strongly coexisting communities are able to supply high levels of overall functioning.

At the pairwise level, coexistence arises from a combination of two mechanisms: niche differences and fitness differences^34^. We tested the relative contribution of these two mechanisms not within a pairwise context but within our three species communities and found that, in line with our expectations, niche differences promoted multifunctionality. However, they did so in interaction with fitness differences, so that the highest overall and net effect multifunctionality occurred when niche differences were high and fitness differences were low (Fig. 3a and S7a). This combination of high niche, and low fitness, differences is also where minimum distance to exclusion is highest^15^. Niche differences might promote high multifunctionality due to complementary effects of niche differentiated species on individual functions. For example, niche differences promoted higher root biomass in the three species communities (Fig. S8a), which could be due to species having complementary rooting depths^35,36^. However, as most individual functions were not impacted by niche differences, it is also possible that species differing in their niches supplied different functions and this variation in the functions supplied by different species increased multifunctionality^27^. However, the interaction between niche and fitness differences also shows another route to high overall and net effect multifunctionality. As well as being high in strongly coexisting communities, multifunctionality was also high in communities with small niche differences and large fitness differences (Fig. 3b and S7b). The presence of such strong competitors in these communities could promote multifunctionality through a type of multifunctional selection effect^7^, i.e., if the strong competitors promote many more functions than the others^37^. Although the ultimate mechanisms driving the niche and fitness differences in our system are unknown, they are likely to arise from multiple causes such as differences in plant physiology^38^, resource use^39^, phenology^40^, and natural enemies^41^. Obtaining all this detailed knowledge would be desirable to explain why these coexistence mechanisms promote multifunctionality. However, our results show another sort of mechanistic understanding. They show that multifunctionality can be predicted from high level mechanisms^42^ such as niche and fitness differences, derived from species intrinsic growth rates and intra and interspecific competitive effects. Therefore, our work establishes a direct mechanistic connection between population dynamics and the functioning of diverse communities.

**Figure 3:**
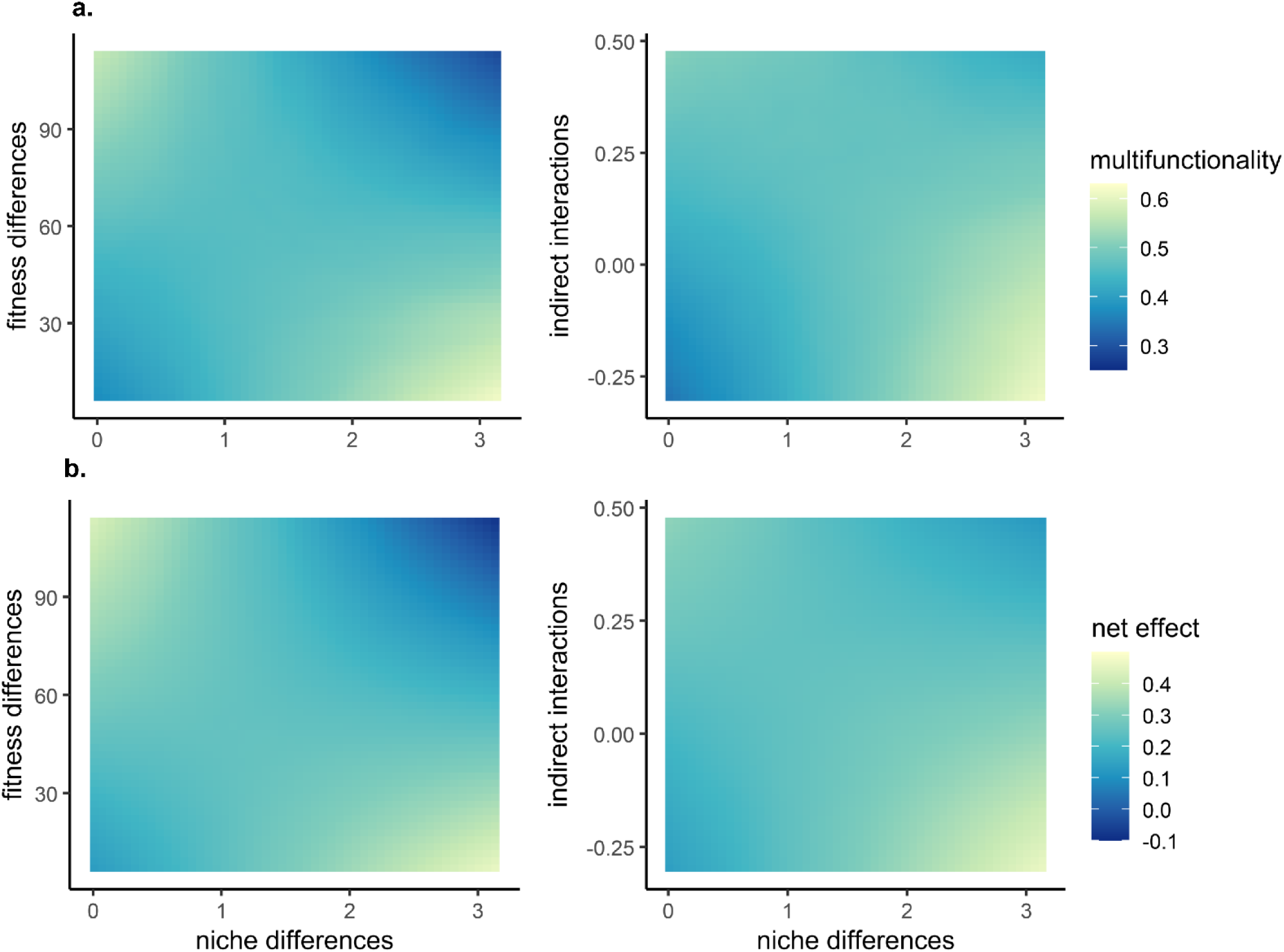
Effects of coexistence mechanisms on multifunctionality and individual functions in control conditions. Panel (a) is a heat map representing the interacting effect of niche differences, fitness differences and indirect interactions on multifunctionality, while (b) represents the interacting effects of these same coexistence mechanisms on net effects (colour = predicted value of multifunctionality or net effect). Niche differences refer to the structural niche differences, fitness differences refer to the structural fitness differences derived with the method from ref. 5.

Our study allowed us to assess the impact of indirect interactions on multifunctionality for the first time. Interestingly, we found that high indirect interactions increased multifunctionality and net effect multifunctionality when niche differences were smaller (Fig 3). These results suggest that high multifunctionality and net effect multifunctionality may also arise within communities where indirect interactions, rather than niche differences, promote coexistence. This may be because indirect interactions can allow species to coexist despite low pairwise niche differences^43,44^, and therefore despite a lack of potential complementarity between them^3^, as is the case for rock-scissors-paper dynamics. However, such types of indirect interactions may have been rare amongst our species as we did not find many possible three species communities where indirect interactions substantially increased coexistence despite small niche differences; instead indirect interactions only seemed to increase coexistence when pairwise niche differences were also fairly high, but insufficient to allow coexistence (at least > 1, Fig. S11b). Indirect interactions may therefore have provided an alternate pathway to high levels of multifunctionality, when structural niche differences were moderate (Fig. 3, Fig. S7). This may suggest that a different set of functions are promoted by indirect interactions vs. niche differences, perhaps because different underlying mechanisms drive multispecies indirect interactions vs. pairwise niche differences, although there is currently very little empirical or theoretical work on this^45,46^. However, we did not observe a clear pattern in terms of some functions being driven more by niche differences or indirect interactions (Fig. S9). For example, we found that root biomass seemed more driven by niche differences while betaglucosidase was reduced by indirect interactions, but on average the individual functions were rarely affected by either coexistence mechanism (Fig. S8). Taken together our results imply that indirect interactions between plants with moderate niche differentiation could drive synergies that lead to several functions increasing overall, and relative to their levels in monocultures. Previous biodiversity-functioning theory has rarely considered how complex plant-plant interactions might translate into ecosystem functioning^47^, but these results suggest that indirect interactions could have important consequences within more diverse communities.

Our experimental approach involved assembling communities that were predicted to differ in the overall strength of coexistence and its underlying mechanisms. Although the interaction coefficients used to derive these predictions were measured in a different experiment, with different soil conditions, and in a different year^4^, the predicted degree of coexistence, and underlying mechanisms, could explain variation in multifunctionality between the newly assembled experimental communities. This suggests that the measured interaction coefficients may be fairly stable and transferable across environmental contexts. This was corroborated by further analyses in which we tested our predictions by using the interaction coefficients from the PaNDiv experiment to simulate the population dynamics of our 12 species in each of the 48 newly assembled communities. Our goal was to determine whether we could predict extinctions occurring in these communities, however, since few extinctions had occurred after two years (only 13 across 60 communities, Table S2), we instead focused on evenness as a metric. We assume that evenness is a more sensitive metric to evaluate short term changes, as species should decline to low levels before going extinct^48^. We therefore fitted models for each of our three species communities, using the interaction coefficients and intrinsic growth rates measured in PaNDiv, and predicted the evenness of the new communities at equilibrium (see methods). These models were able to predict the observed evenness in the communities after two years (Fig. 4). We did not observe as low evenness as predicted by our models (Fig. 4), perhaps because the communities had not had time to fully reassemble and for low performing species to be excluded. However, we feel it is encouraging that our predictions are significantly related to the observed values, given that we used data from a single growing season and very simple population models to estimate the interaction coefficients, which ignore much of the complexity of perennial plant population dynamics^49^. These results indicate that such simple approaches to measuring perennial plant interactions provide valuable information which can predict changes in biodiversity, as well as overall multifunctionality levels.

**Figure 4:**
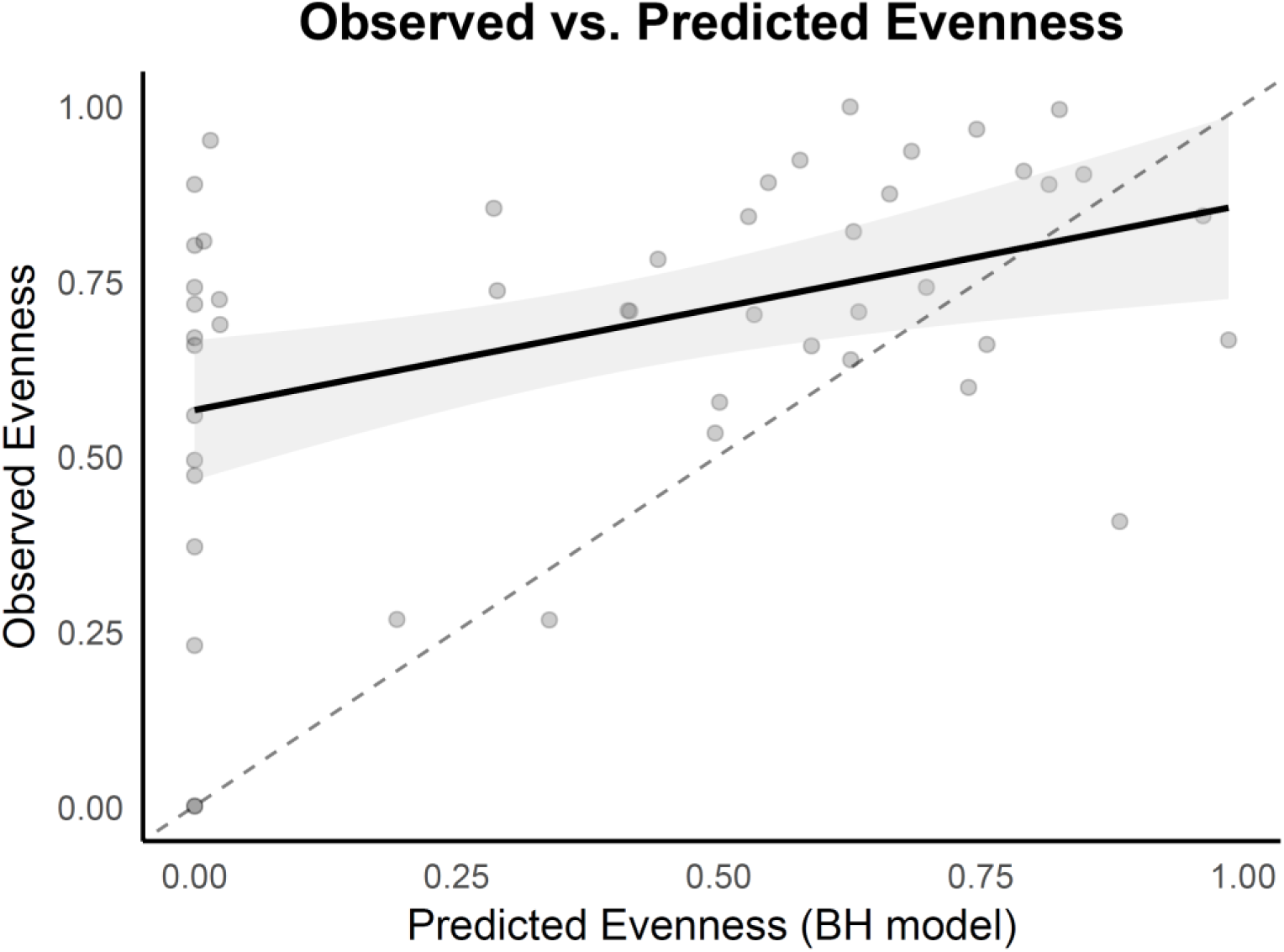
Predictions of evenness with Beverton-Holt (BH) model using interaction coefficients from June 2020 measured in PaNDiv experiment, vs. evenness observed in June 2022 in our triplet experiment (each dot represents a plot).

Multifunctionality is strongly dependent on the number and identity of functions included^24,50^. Here, we included seven above and belowground functions, aiming to represent as many aspects of functioning as possible. Our results show that the number of functions and their ecological compartment had little impact on the link between coexistence and multifunctionality (Fig. S6). However, we observed a lot of variation in effects when only a few functions were included and very few effects on individual functions. Biodiversity effects on functioning often accumulate over time^51,52^, particularly belowground^53^, and effects of coexistence strength might also accumulate over time. Investigating the link between coexistence and functioning over time could therefore determine how the effects of coexistence mechanisms arise from the accumulation of many small effects on individual functions. For instance, increases in biomass might lead to increased carbon input into the soil, leading to increased microbial activity and rates of nutrient cycling and therefore further increases in plant biomass over time^18^. Extending our results to higher diversity levels would also be valuable^11^. In this study we examined the effects of pairwise interactions and indirect effects among three species. Future research could explore how these coexistence mechanisms, especially indirect effects but also higher order interactions^54^, contribute to functioning across and within higher levels of species richness. Some work has suggested that higher order interactions might drive more positive effects of biodiversity on biomass, by reducing interspecific competition at higher diversity^55^, and it would be interesting to extend this work to effects on multifunctionality. Further experiments assembling communities that vary in the predicted extent of indirect and higher order interactions would be very valuable.

We provide the first experimental evidence for a connection between coexistence mechanisms and multifunctionality. The finding that more strongly coexisting communities provide more functions at high levels shows that the fundamental population processes regulating biodiversity also scale up to impacts on ecosystem functioning. As biodiversity loss arises from a disruption of coexistence mechanisms, our results suggest that by altering species interactions, global change drivers might reduce multifunctionality, even before biodiversity is lost. Niche differences and indirect interactions represent different pathways by which species interactions stabilize communities and promote coexistence, and our results showed that they also provided alternative routes to high multifunctionality. Revealing these effects required a new experimental approach and this could be extended to investigate the physiological, morphological, and phenological mechanisms affecting the drivers of multispecies coexistence mechanisms. Assembling communities predicted to differ in coexistence strength and its underlying mechanisms has the potential to validate model predictions by examining the development of the communities and the link between coexistence and ecosystem functioning and stability. Our results show that the conditions for stable coexistence and high functioning are related, which should allow a more predictive and mechanistic link between changes in diversity and ecosystem functioning.

## Material and Methods

### Structural coexistence

We set up the experiment manipulating coexistence mechanisms, using results obtained in the PaNDiv experiment located in Münchenbuchsee, Switzerland. PaNDiv manipulates plant species richness, with plots sown with 1, 4, 8, or 20 species (the experiment also involves nitrogen and fungicide treatments but these are not considered here). In September 2019, phytometers of 18 perennial plant species were planted into different competition neighbourhoods with the 2m x 2m plots, to estimate interaction coefficients. We planted 10 individuals of each species into its respective monoculture to estimate intraspecific competition coefficients (αii). Then, we planted 3 replicates of every possible pairwise combination of species in heterospecific plots to estimate every interspecific competition coefficient (αij). Finally, we planted 4 replicates of each species growing alone in plots where we prevented the growth of competitors by covering the plot with landscape fabric. We used these phytometers to estimate λi, the intrinsic growth rate for every species. In May and August 2020, we sampled the biomass of the phytometers, and we recorded the percentage cover of every neighbour within a circle of 20 cm radius. In order to compute all values of α and λ, we ran a negative binomial regression model for each focal species (“GLMMTMB” package, Brooks et al. 2017, ref. 56) where the response variable was the dry weight of the phytometer (in mg.). Biomass was predicted by the cover of each of the 18 individual species (separately assessed, including conspecifics) and interactions of these with each of PaNDiv experiment treatments. The model was run on data from the two sampling periods (June and August) in order to get a mean interaction value for each pairwise species combination, and mean intrinsic growth rate values for each species^4^.

Based on these intrinsic growth rates and competition coefficients, structural coexistence was computed for all possible combinations of three species. For each three species community we calculated the structural niche difference, intrinsic growth rate ratio (structural fitness differences), overall strength of indirect interactions and coexistence feasibility^5^. We also computed the minimum distance to exclusion for the three species communities, as the time for the first species to be excluded, following the method from Allen-Perkins et al. 2023 (Supplementary Method 1, Fig. S10, ref. 15).

### Experimental design

Using these results, we selected 48 three species communities that maximised variation in niche differences and indirect interactions, while also minimising the correlation between the two (Fig. S11b). We focussed on niche and indirect interactions as variation in these was shown to be most important in determining whether a three species community could coexist. Some of the communities predicted to coexist have higher fitness differences than niche differences, but can still coexist thanks to average indirect interactions, while others are able to coexist in a more classic way, thanks to large niche differentiation. Our selection resulted in communities with a subset of 12 plant species and we tried to limit as much as possible the over or under representation of every species in the three species communities, and ensured that the presence of a particular plant species was not correlated with niche differences or indirect interactions. We also grew all 12 species in monocultures for a total of 60 plots (see plot species composition in Table S1).

We sowed the communities in May 2021, in outdoor 1 x 1 m plots, watered them every week as long as they were in the seedling stage and monitored species establishment (percentage cover) during the first year. We sowed enough seeds to get a density of 1000 individuals per 1m^2^ plot, which is the same starting density of individuals as in the PaNDiv experiment. We sowed seeds of each species in equal proportion, corrected for germination rates. We used a compilation of germination rate data previously obtained from literature or from our own germination experiments. One species, *Poa trivialis*, had to be resown in September 2021 because it didn’t establish well. Plots were weeded three times in April, July and September, at the same time as the PaNDiv experiment.

### Ecosystem functions

In order to link coexistence mechanisms with ecosystem multifunctionality, we measured a set of seven functions. Three of them were linked to aboveground processes: plant shoot biomass, leaf fungal pathogen and herbivory damage, and four were proxies of belowground processes: root biomass, litter decomposition, beta-glucosidase and acid phosphatase activity. Plant shoot biomass was sampled by cutting the total biomass of each plot, leaving a minimum of 5 cm of aboveground material, drying it at 80°C for three days and weighing it to record the dry weight (in g.). Damage was recorded by randomly selecting 10 leaves of different adult individuals for each species in the plot, measuring the proportion of their leaves damaged by herbivores and fungi, as well as visually assessing the average percentage cover of each type of damage in each leaf. We then calculated the community weighed mean (CWM) of the damage using the plant percentage cover visually recorded in the plot in May 2022 as: 𝐶𝑊𝑀 = ∑ 𝑝_𝑖_ ∗ 𝑛_𝑖_, with 𝑝_𝑖_ the percentage cover of each species within the plot, and 𝑛_𝑖_ the mean damage recorded for this species in the plot. Root biomass was sampled by coring two portions of the plot, mixing the two samples together, weighing them and sieving them to separate the root from the soil. Fresh roots were then dried at 80°C for two days, weighed, and root biomass was calculated as the amount of dry root mass (in mg) for a g of soil. Litter decomposition was sampled by filling two litter bags per plot with around 10g of litter. Each bag was of 5mm mesh size on the top and 0.2mm on the surface in contact with the soil. The litter used was from the same plot, sampled during the most recent biomass harvest (August 2022). The bags were left to decompose on top of the bare ground soil for two months and were harvested, dried in the oven at 80°C for three days and weighed. The litter bags were the same as those used to measure litter decomposition on PaNDiv in Pichon et al. 2020^57^. Soil enzymatic activity was measured by coring two samples in the middle of each plot, sieving them and freezing them in order to limit the chemical release from microbes. We tested the concentration of beta-glucosidase with p-nitrophenyl-D-glucopyranoside, and the concentration of phosphatase with p-nitrophenyl-phosphate, along with a calibration curve constructed for each enzyme, following the protocol from Tabatabai 1994^58^. Functions were sampled between May 2022 and March 2023. For biomass, measured twice during this time period, we calculated the mean value for each plot.

### Analyses

#### - Multifunctionality

To calculate multifunctionality, we first checked that there was no strong correlation between each of the seven functions included in the study. We then checked that functions followed an approximately normal distribution, as highly skewed distributions can lead to high or low influence of a given function on multifunctionality (Pichon et al. 2024, ref. 7), and we log-transformed the leaf fungal pathogen and herbivory damage data to get closer to normality. Multifunctionality was then assessed for multiple thresholds^23,24^. We scaled each function per plot to the 5 highest values found across plots, in order to reduce the impact of outliers, and then calculated the proportion of functions that exceeded each threshold from 0.5 to 0.8 by steps of 0.05. After transformation, no particularly strong correlation was found between any individual functions and multifunctionality (ranging across all thresholds, in absolute values, from 0.01 to 0.63 Fig. S12), suggesting that no one function drives the response.

In order to assess the link between coexistence mechanisms and overall multifunctionality, we then analysed all the multifunctionality values from different thresholds at the same time^7^. This allows us to calculate the average effect of coexistence mechanisms across multifunctionality thresholds. In these models, we also need to account for the potential effect of individual species, especially since we could not include all combinations of three species in our experimental design. For this, we fitted linear models with multimembership random effects using a presence/absence matrix of species as a random factor. Using this method, we correct for individual species effects and the greater similarity that exists between plots sharing more species. These models contain multiple measures of multifunctionality per plot and although we do not include a specific plot random effect, the multimembership random effect corrected for the plot effect because all measures from the same plot are counted as being maximally similar. However, they also score plots sharing more species as being more similar and therefore simultaneously correct for spatial pseudo replication and pseudo replication due to shared species between plots. It was not possible to include a second random effect for multifunctionality threshold (categorical) in these multimembership models, as in Pichon et al. (2024). However, we also ran alternative mixed models without the correction for similarity in species combinations (i.e., without a multimembership random effect) but with multifunctionality thresholds as a categorical random factor, and the variance of this random factor was very close to 0, showing that species identity was a better random factor to use in this study.

We ran two types of multimembership models. First, we included the minimum distance to exclusion as a continuous variable representing the degree of stable coexistence in the community. Then, we ran a model including all coexistence variables, i.e. structural niche differences, intrinsic growth rate ratio and indirect interactions. We included in the model the interaction between structural niche differences and intrinsic growth rate ratio as well as the interaction between structural niche differences and indirect interactions. For each model we also included the multifunctionality threshold as a continuous variable and tested its interaction with all of the explanatory variables included.

#### - Net effects

In addition to the overall measurement of multifunctionality, we also include a new multifunctionality measure that quantifies the extent to which our three species communities provide higher multifunctionality than their constituent monocultures, net effect multifunctionality. In order to calculate net effect multifunctionality^25^, we calculated for each species in each plot, the predicted value of each function, using the function values sampled in the monocultures. We then calculated the predicted values of each function in the three species communities, as the average of the three values per function (i.e., assuming the species abundance to be equal to the sown abundance). We then calculated the net effect as the difference between observed and predicted values for each individual functions in each plot. Net multifunctionality was calculated using the same method as overall multifunctionality and was analysed with similar multimembership models.

#### - Individual functions

We also assessed the impact of overall coexistence, structural niche, structural fitness differences and indirect interactions on each individual function we sampled. Similar to overall and net multifunctionality, we ran two multimembership models for each function, separately assessing the effect of a continuous variable of coexistence, minimum distance to exclusion, then assessing the effect of the three separate components of coexistence, while correcting for individual species effect, including them as a random factor in the form of a presence/absence matrix.

#### Assessing the impact of function number and combination on overall and net effect multifunctionality

In order to verify how the number and identity of functions influenced the effect of minimum distance to exclusion on overall and net effect multifunctionality, we performed an analysis following Meyer et al. 2018 (ref. 33), where we computed all possible combinations of functions between 2 and 7 functions (n = 120 possible combination of functions) and we calculated the multifunctionality value for each plot and each threshold (from 0.4 to 0.8 thresholds by steps of 0.05, n = 103’680 data points in total). A similar type of analysis can also be found in ref. 7 and 57. Then, we ran the multimembership models for each of these combinations and extracted every regression coefficient of the fixed effect of the minimum distance to exclusion on both the multifunctionality (Fig. S6a) and the net effect multifunctionality (Fig. S6b).

#### - Evenness predictions

Finally, we wanted to compare the abundance shifts of our communities with the predicted abundance shifts using the interaction coefficients sampled on the PaNDiv experiment. For this, we fitted a Beverton-Holt model of population dynamics following this equation:

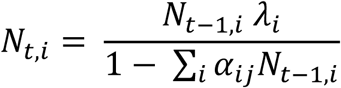

Where 𝑁_𝑡,𝑖_ is the abundance of species i at time t, 𝜆_𝑖_ is the intrinsic growth rate of species i, 𝛼_𝑖𝑗_ is the competition coefficient between i and neighbour species j (for i = j, 𝛼_𝑖𝑗_ = 𝛼_𝑖𝑖_ is the intraspecific competition coefficient). To parametrise this model, we used the interaction coefficients between our 12 species sampled in June 2022, with all species at equal initial abundances 𝑁_0_ (as it was sown in the beginning of the experiment). We derived the predicted Pielou’s evenness^60^ from this model, at a time point we considered long enough to approach a realistic equilibrium (15 years). We then calculated the evenness of our communities in June 2022. We chose to focus on coefficients and observed evenness in June, since our estimates of June 2020 coefficients from the PaNDiv experiment are more robust than the coefficients from August 2020. This is because the experiment is cut in June and many phytometers were lost or dead following the cut, which reduced the sample size in August 2020^4^. We then correlated predicted evenness with observed evenness in June 2022.

## Data availability

All data and code that support the findings of this study are available in a public GitHub repository at this address: https://github.com/cardaips/structural_coexistence_multifunctionality. A dryad repository containing data linked with this study can be found under the identifier DOI: 10.5061/dryad.3ffbg79zh.

## Acknowledgments

We are grateful to the whole PaNDiv team, especially Sylvain Chartier, Mervi Laitinen, Hugo Vincent and all the helpers that worked on the maintenance of both the coexistence and multifunctionality experiments. We also thank Seraina Cappelli for her role in setting up the PaNDiv experiment. We are also grateful for the students that participated in the data collection of ecosystem functions: Jerome Bechtold, Eva Anna Burgunder, Silja Eller and Valentina Hurni Mendoza, and the team that participated in collecting the cover data within the PaNDiv experiment: Vera Alessandrello, Hannah Bratschi, Eli Bucher, Tala Bürki, Géraldine Chavey, Chiara Durrer, Matthieu Gauvrit, Benjamin Herren, Fabian Heussler, Vinciane Horner, Sandy Kalaydjian, Nadia Maaroufi, Olivier Magnin, Dmitry Maryasov, Anja Michel, Thu Zar Nwe, Barryette Oberholzer, Scarlett Peréz Gordillo, Valentin Pulver, Georges Saumier, Nynke Van Duijin, Joseph Volery and Lia Zehnder. PaNDiv was funded by the Swiss National Science Foundation (Awards 31003A_160212, 310030_185260 and 310030_215442). Oscar Godoy acknowledges financial support provided by the Spanish Ministry of Science, Innovation and Universities (MICIU) and by the European Social Fund through the TASTE (PID2021-127607OB-I00) and BIOTA (EUR2023-143472) projects.

## Author contributions

E.A. and O.G. designed the coexistence experiment carried out within the PaNDiv experiment to extract competition coefficients and intrinsic growth rates for the 12 plant species. C.D., E.A. and O.G. designed the multifunctionality experiment. C.D. sampled the data from the coexistence experiment, C.D. and M.G. sampled the ecosystem functioning data within the multifunctionality experiment. C.D., E.A, O.G. and N.P. contributed to the data analyses. C.D., E.A. and O.G. wrote the first draft of the paper, all co-authors contributed extensively to its subsequent revisions.

