## Supplementary material for "Stably coexisting communities deliver higher ecosystem multifunctionality"

- Supplementary Method 1: Structural stability versus Modern Coexistence Theory (MCT) frameworks

The structural stability framework[5] is an expansion of the MCT framework[34] from the pairwise level to an arbitrary (*n*) number of species, using a translation in the form of a geometrical projection of each mechanism of coexistence.

Similarly to the MCT, the mathematical base of the structural stability framework is (1) the matrix of interactions between *n* number of species, and (2) each species intrinsic growth rate. From this base, MCT typically derives two mechanisms at the pairwise level: the stabilising mechanisms (also called niche differences) and the equalising mechanisms (i.e., competitive ability differences or fitness differences). Both of those processes can be compared to one another to assess whether the two species can stably coexist or not (here, coexistence is binary outcome, competitive exclusion or coexistence).

On the other hand, structural stability expands this MCT framework, by building a geometric space from the interaction coefficients (the feasibility domain), where a community of species can coexist beyond the pairwise level. This framework also allows us to consider all signs of interactions (including facilitative ones), whereas MCT can only comprise competitive ones.

At the pairwise level, the feasibility domain computed by the structural stability approach is entirely determined by the niche differences between two species. Beyond the pairwise levels however, the size of the feasibility domain is also modified by indirect interactions that emerge from species-rich environments. As such, indirect interactions might promote opportunities for coexistence (enlarge the feasibility domain) or reduce them (reduce the feasibility domain).

On the contrary, the fitness differences computed by the structural stability framework are entirely determined by the ratio of *n* species growth rates. Unlike the MCT framework, they don't comprise species response to competition in the equalising mechanisms. As a result, the feasibility domain is entirely dependent on the matrix of interactions, which breaks the mathematical correlation between niche and fitness differences that was present in the MCT framework.

If the vector projected from the ratio of intrinsic growth rates falls inside the feasibility domain, then the community of species stably coexists. This geometrical approach also led to the development of a metric describing coexistence in a more continuous way, as the minimum amount of disturbance that the community is able to undergo before one of the species goes extinct (minimum distance to exclusion[15], Fig. S10).

- Table S1: The plot species composition in Ostermundigen, consisting of 12 monocultures of common perennial grassland species: *Bromus erectus (Be)*, *Crepis biennis (Cb)*, *Daucus carota (Dc)*, *Festuca rubra (Fr)*, *Holcus lanatus (Hl)*, *Lolium perenne (Lp)*, *Plantago media (Pm)*, *Poa trivialis (Pt)*, *Prunella grandiflora (Pg)*, *Rumex acetosa (Ra)*, *Salvia pratensis (Sp)*, *Taraxacum officinale (To)*, and 48 plots of three species combinations of those species.

| plot n° | Species name | | | plot n° | Species name | | |
|---|---|---|---|---|---|---|---|
| 1 | To | Dc | Ra | 31 | Pt | Pg | Ra |
| 2 | Cb | Dc | Pg | 32 | Be | Pt | Sp |
| 3 | Fr | Hl | Ra | 33 | Be | Sp | Ra |
| 4 | Lp | Pt | Dc | 34 | Fr | Pt | Sp |
| 5 | Be | Hl | Dc | 35 | Hl | Pt | Ra |
| 6 | Fr | To | Ra | 36 | Fr | To | Dc |
| 7 | Lp | Cb | Sp | 37 | Lp | Cb | Pg |
| 8 | Pt | Dc | Sp | 38 | Hl | Pg | Pm |
| 9 | Fr | Pt | Pm | 39 | Be | Fr | To |
| 10 | Be | To | Dc | 40 | Cb | Ra | Pm |
| 11 | Pt | Cb | Sp | 41 | Fr | Pt | Cb |
| 12 | Be | Fr | Dc | 42 | Be | Hl | Pt |
| 13 | Be | Hl | To | 43 | Lp | Cb | Dc |
| 14 | Sp | Ra | Pm | 44 | Lp | Cb | Ra |
| 15 | Be | Pg | Ra | 45 | Fr | Hl | Sp |
| 16 | Be | Pt | Pm | 46 | Be | Lp | Dc |
| 17 | Be | Pt | To | 47 | Fr | Lp | Pt |
| 18 | Be | Cb | Ra | 48 | Pg | Ra | Pm |
| 19 | Cb | To | Ra | 49 | Pg | | |
| 20 | Pt | Dc | Pg | 50 | Sp | | |
| 21 | Pt | Cb | Ra | 51 | Cb | | |
| 22 | Cb | To | Pm | 52 | Dc | | |
| 23 | Lp | To | Ra | 53 | Fr | | |
| 24 | Be | Pt | Pg | 54 | Hl | | |
| 25 | Be | Ra | Pm | 55 | Lp | | |
| 26 | Fr | Hl | To | 56 | Pm | | |
| 27 | Hl | Sp | Pg | 57 | Pt | | |
| 28 | Fr | Hl | Dc | 58 | Be | | |
| 29 | Hl | Lp | Ra | 59 | Ra | | |
| 30 | Fr | Ra | Pm | 60 | To | | |

- Table S2: List of extinctions that occurred in the plots between 2021 and 2022. Species abbreviation list can be found in Table S1.

| **species extinct** | **plot n°** |
|---|---|
| Dc | 2 |
| Pt | 9 |
| To | 10 |
| Pt | 16 |
| To | 19 |
| Dc | 20 |
| Pt | 24 |
| Pt | 31 |
| Pt | 32 |
| Pt | 34 |
| Pt | 41 |
| Hl | 42 |
| Pm | 48 |
| **Total number of extinctions** | n = 13 |

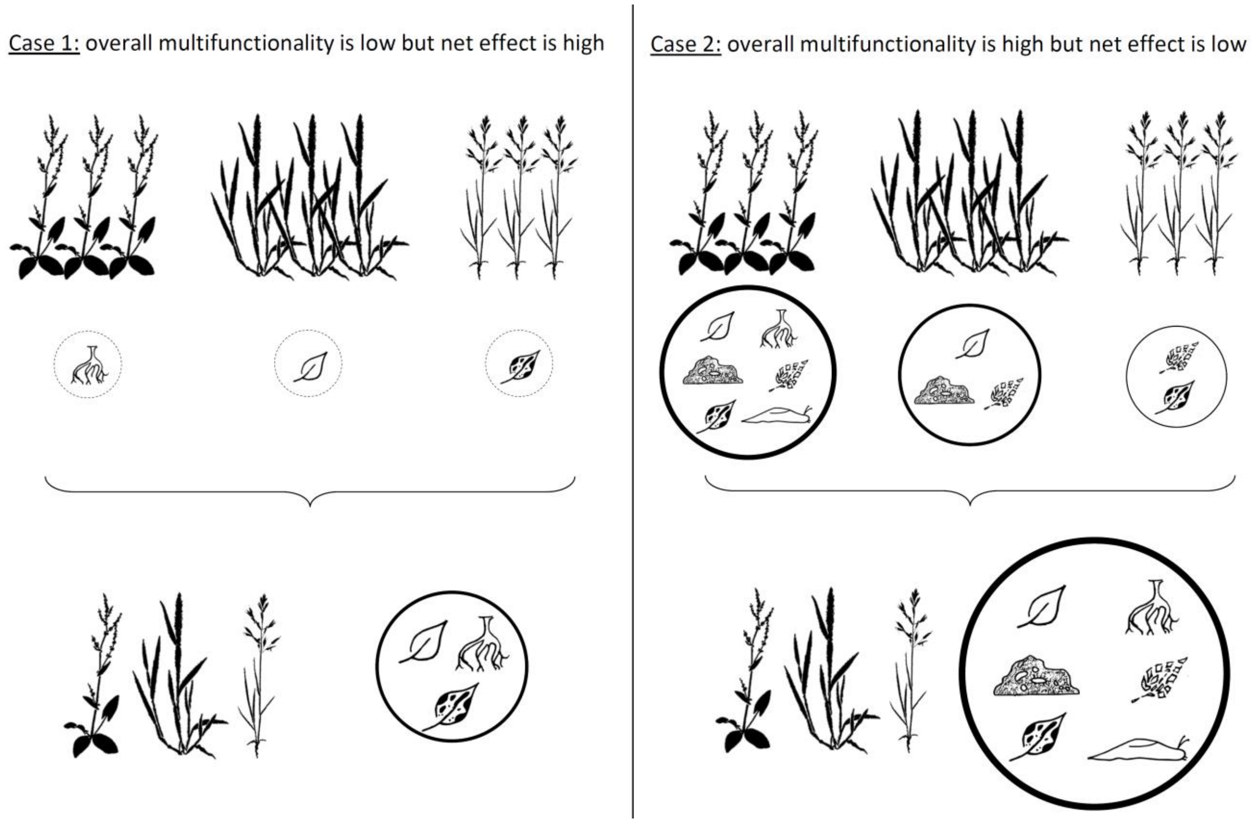


- Figure S1: Two scenarios where overall effects and net effects of multifunctionality are decorrelated: in circles are the functions provided at high levels by each community (monospecific or heterospecific), the outline and size of the circles indicate the overall level of multifunctionality, while the difference between three-species communities and monocultures constitute the net effect.

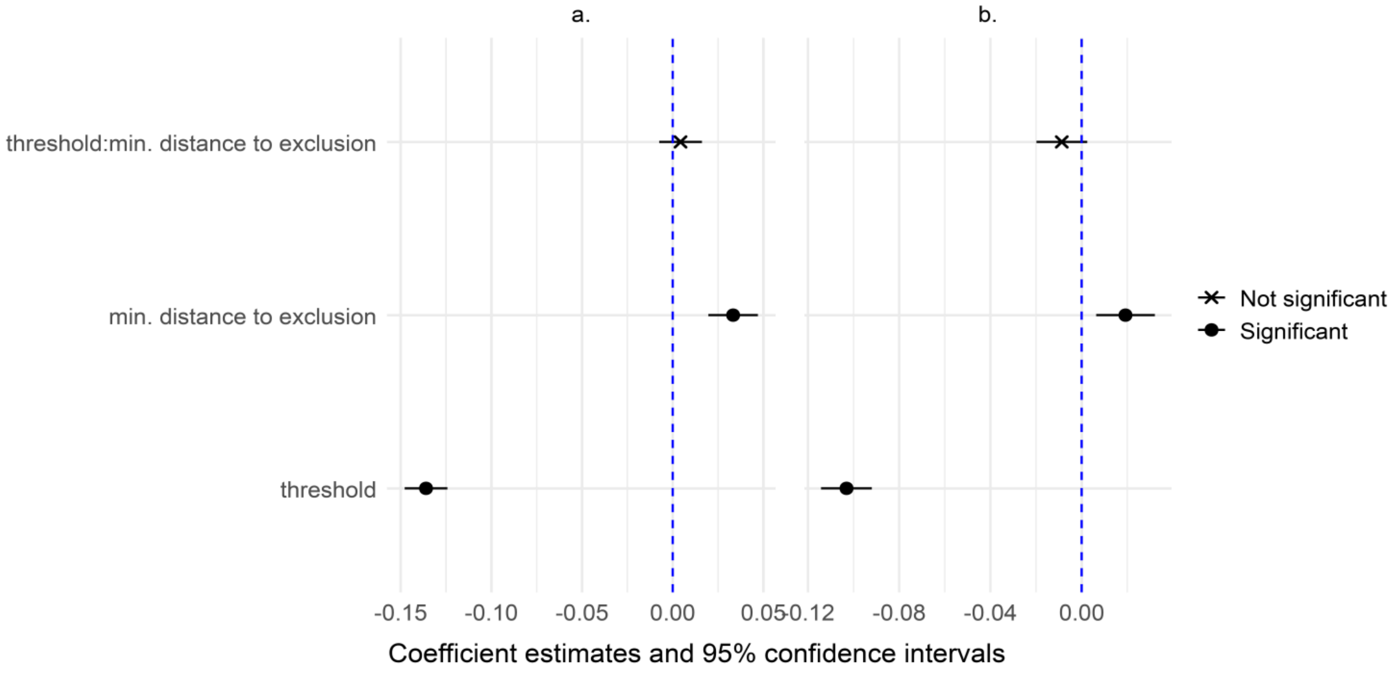


- Figure S2: Multimembership model outputs of overall coexistence impact on multifunctionality (a) and net effect (b) of multifunctionality. These model outputs represent the mean effect and confidence intervals of how multifunctionality obtained within our 3 species communities is impacted by the minimum distance to exclusion of one of the species. threshold = multifunctionality threshold (as a continuous fixed effect).

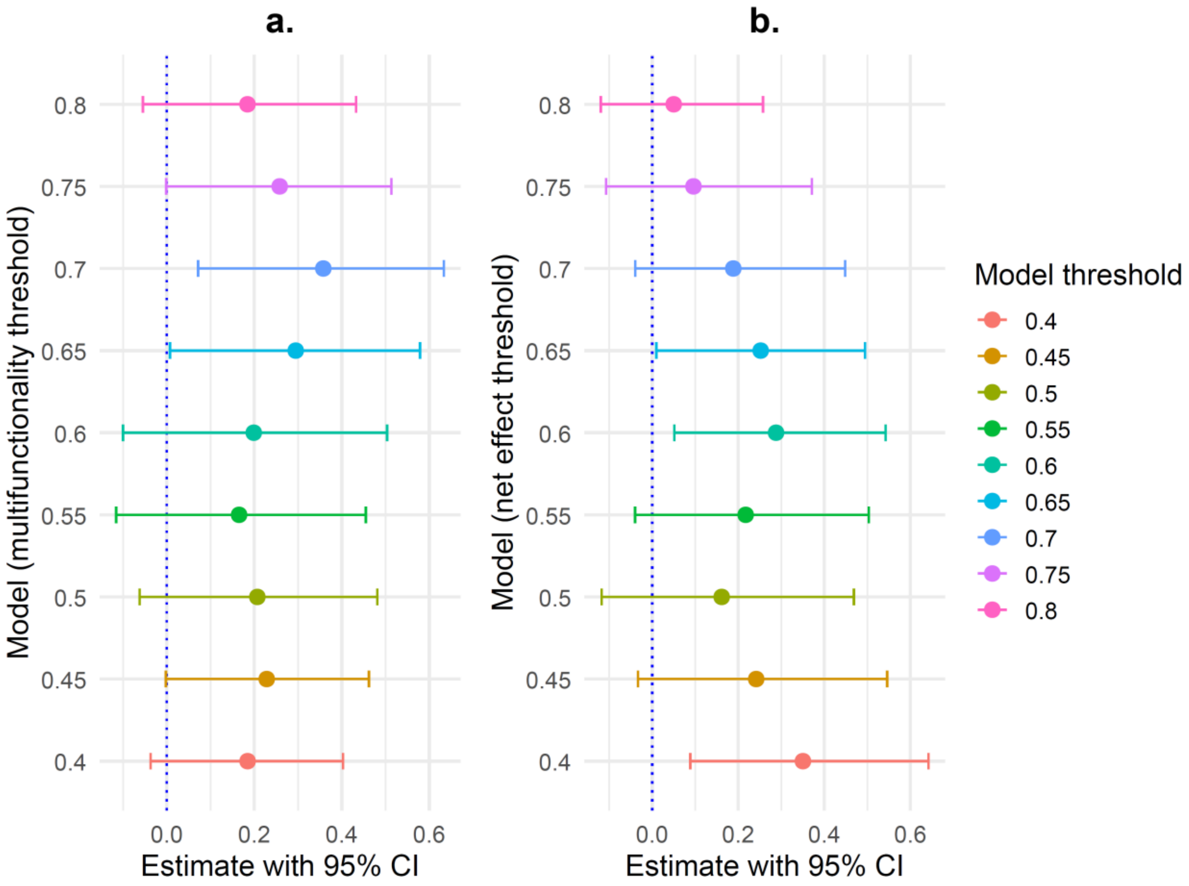


- Figure S3: Effect of minimum distance to exclusion on (a) multifunctionality and (b) net effects for the 9 thresholds, individually. For each threshold, a model with multimembership random factor was used and the estimate and standard error was computed. The global effect of minimum distance to exclusion using all threshold combined in the same dataset can be found in Fig. S2.

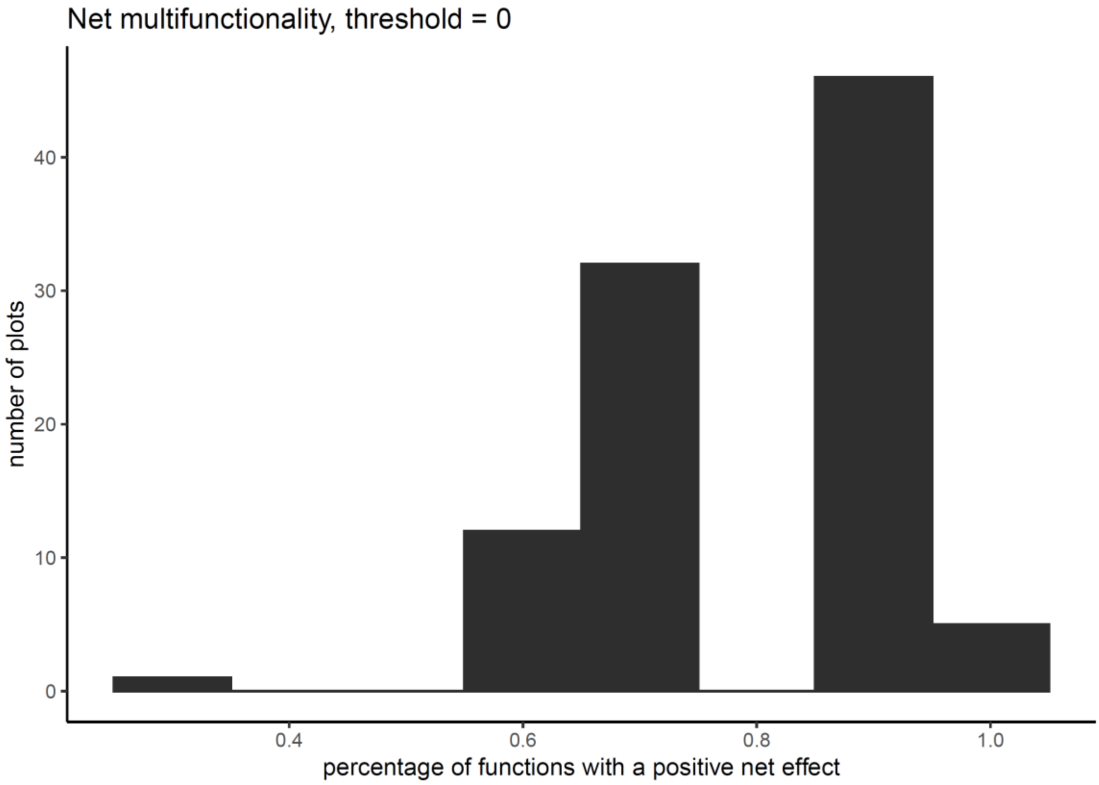


- Figure S4: Histogram of the percentage of functions surpassing their predicted values of functioning in three species plots compared to their respective monoculture.

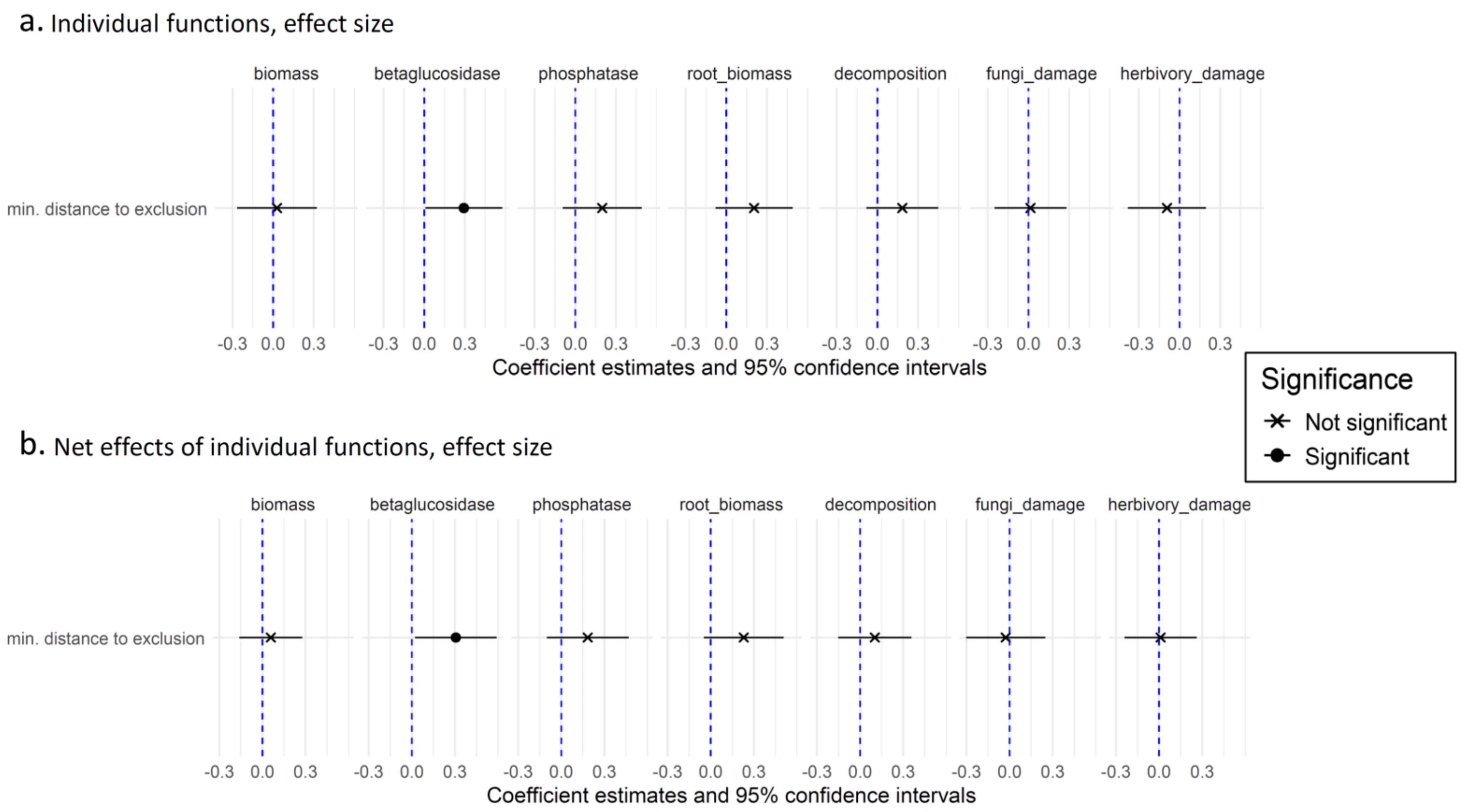


- Figure S5: Multimembership models output of overall coexistence impact on (a) individual functions and (b) their respective net effects. Each panel describes a function measured on our 3 species communities (and their respective monocultures for net effects), and how minimum distance to exclusion (min. distance to exclusion) of one of the species impacts the level of this function.

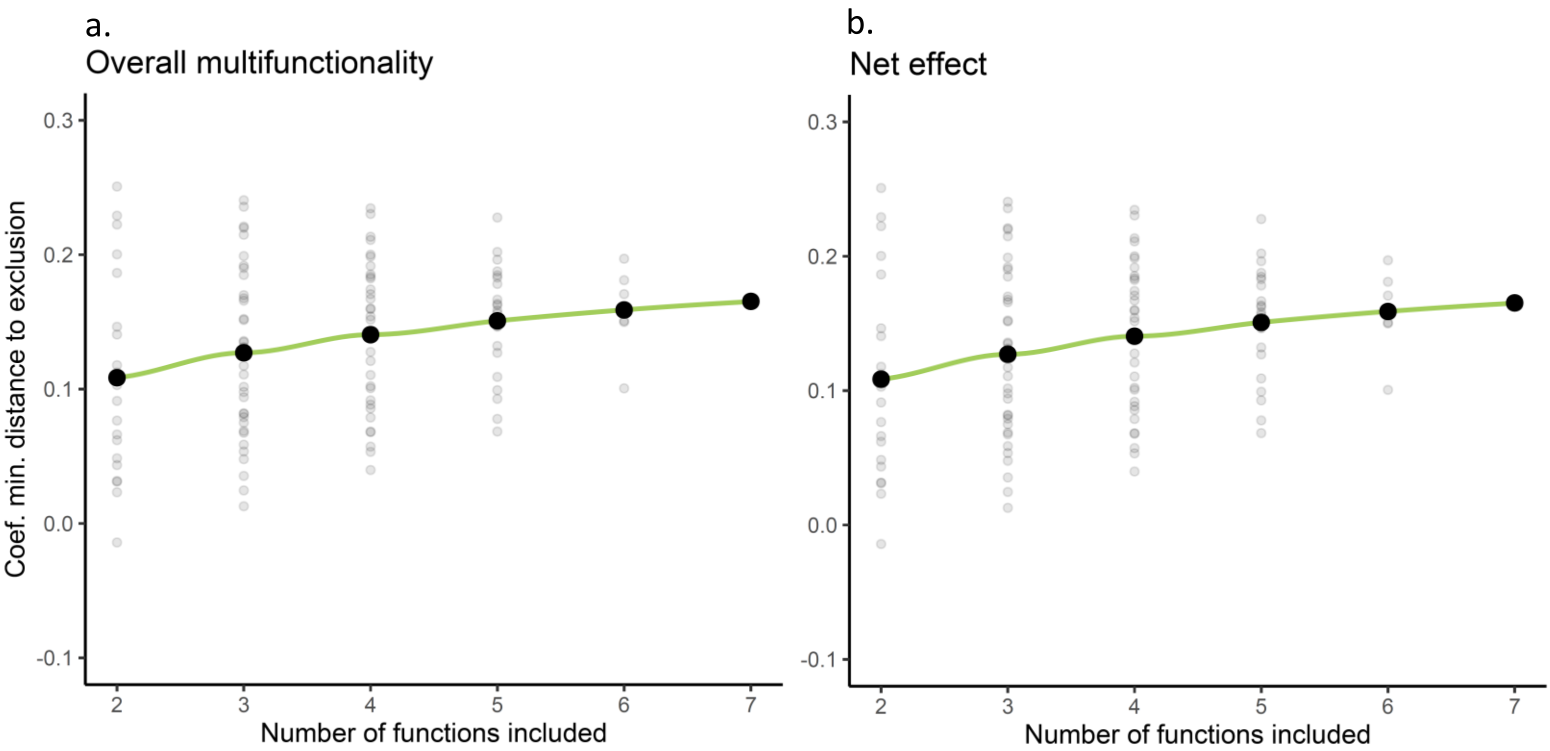


- Figure S6: Regression coefficient of the minimum distance to exclusion impact on (a) the multifunctionality and (b) the net effect, for all possible combinations of individual functions, from 2 to a total of 7 functions. Grey dots are the fixed effect coefficients (n = 120) and the green curve and black dots represent the mean values for each combination level.

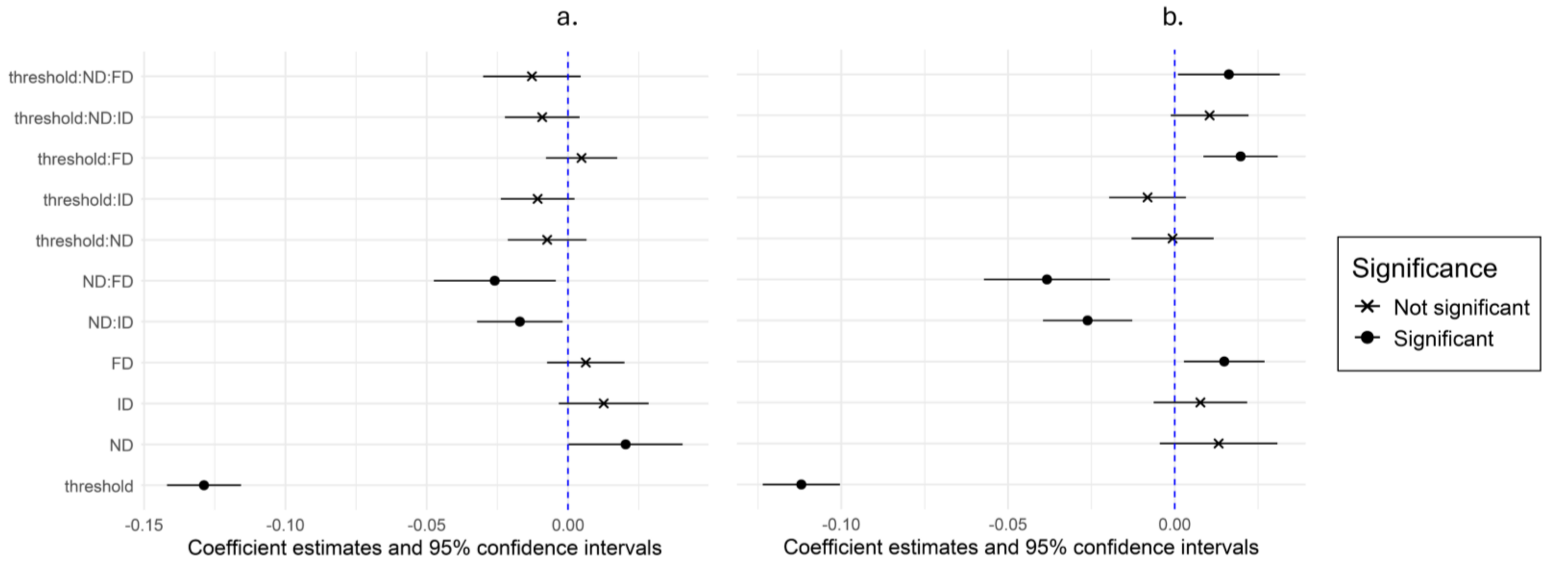


- Figure S7: Multimembership model outputs of each coexistence mechanisms impact on multifunctionality (a) and net effect (b) of multifunctionality. These model outputs represent the mean effect and confidence intervals of how multifunctionality obtained within our 3 species communities is impacted by niche difference, fitness differences, indirect interactions and their interactions. ND = niche differences, FD = fitness differences, ID = indirect interactions, threshold = multifunctionality threshold (as a continuous fixed effect).

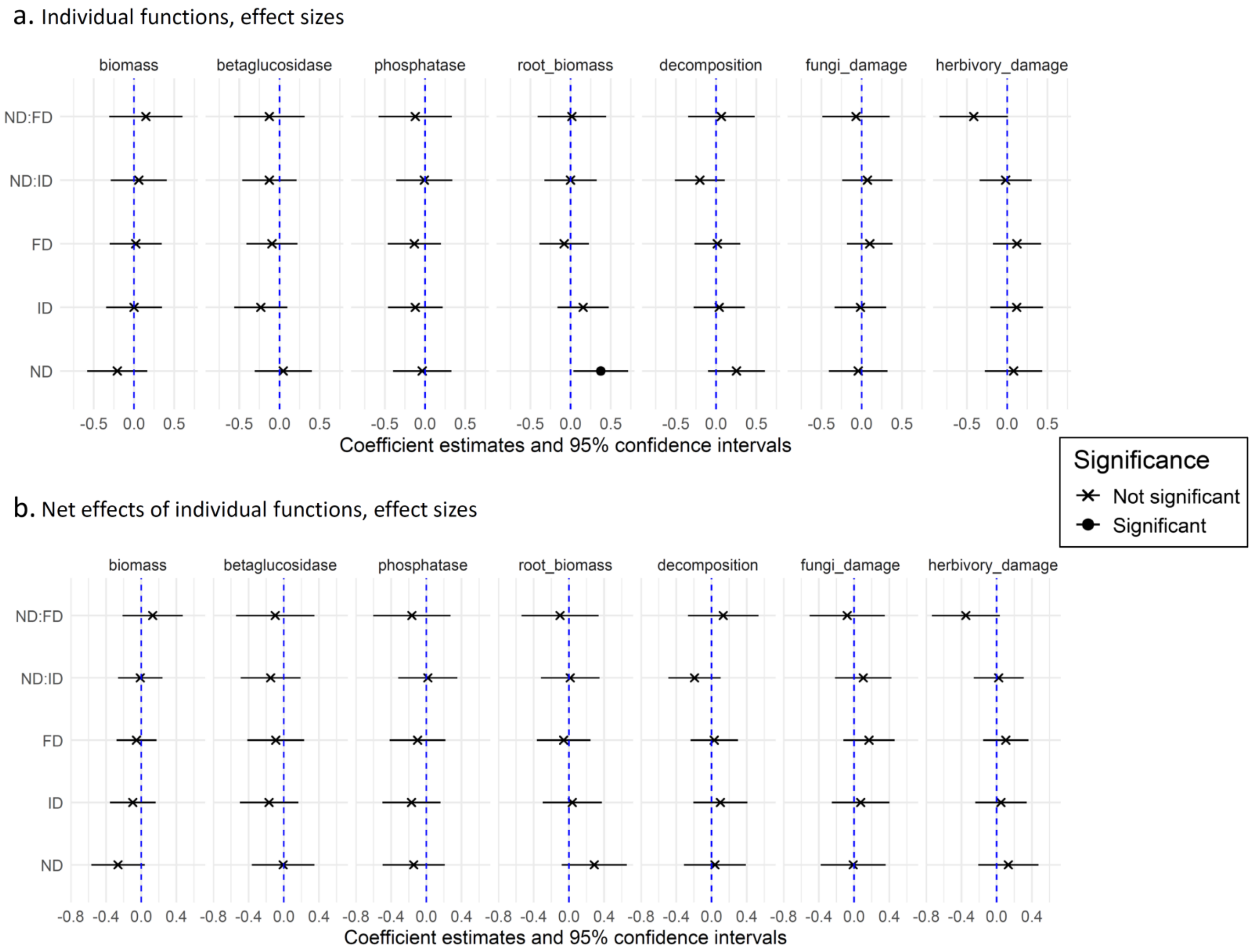


- Figure S8: Multimembership models output of the impact of each coexistence mechanism on (a) individual functions and (b) their respective net effects. Each panel represents a function measured on our 3 species communities (and their respective monocultures for net effects), and how niche differences, fitness differences and indirect interactions of one of the species impacts individually or in synergy the level of this function. ND = niche differences, FD = fitness differences, ID = indirect interactions.

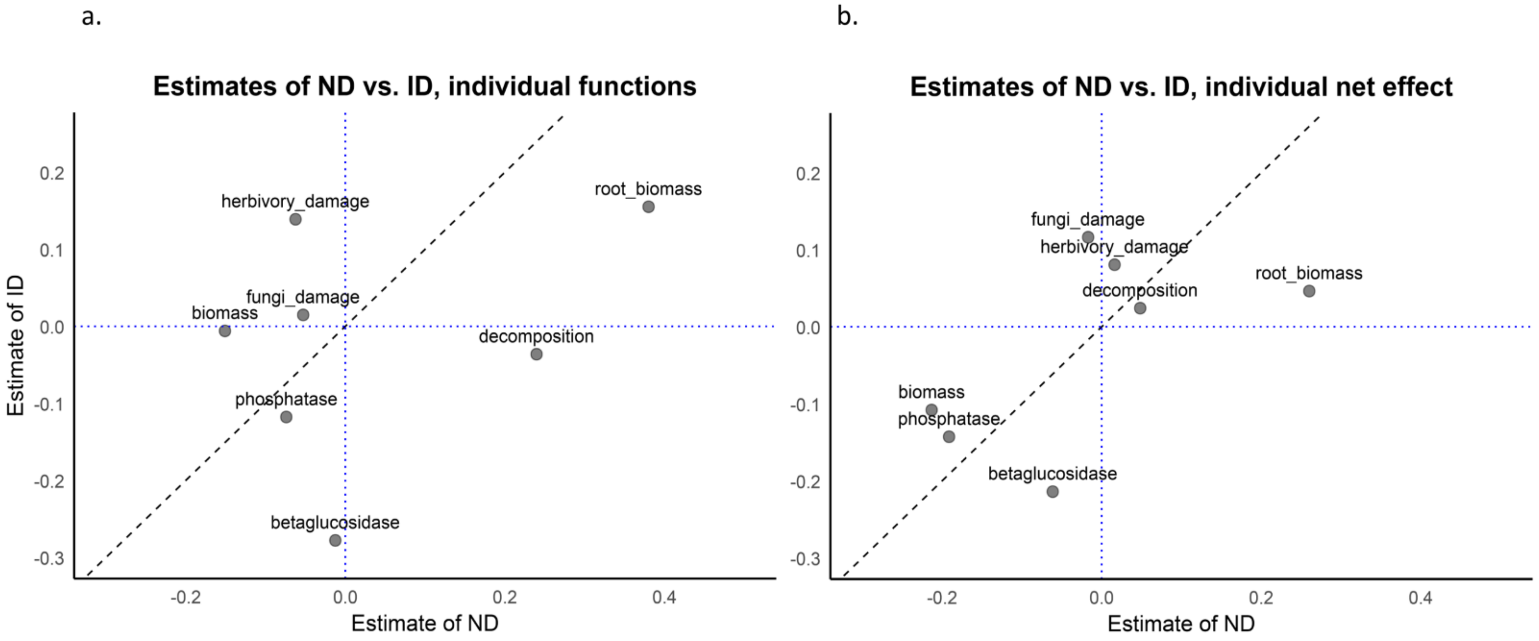


- Figure S9: Estimates extracted from multimembership models of niche differences versus indirect interactions effects on (a) each individual functions and (b) their respective net effect. The 1:1 line in black represents the cases where niche differences and indirect interactions impact functioning to the same extent (when they are of the same sign). If a dot is close to the horizontal dotted line in blue, then the effect of indirect interactions on this function is weaker, similarly if a dot is close to the vertical dotted line in blue, then the effect of niche differences on this function is weaker. ND = niche differences, ID = indirect interactions.

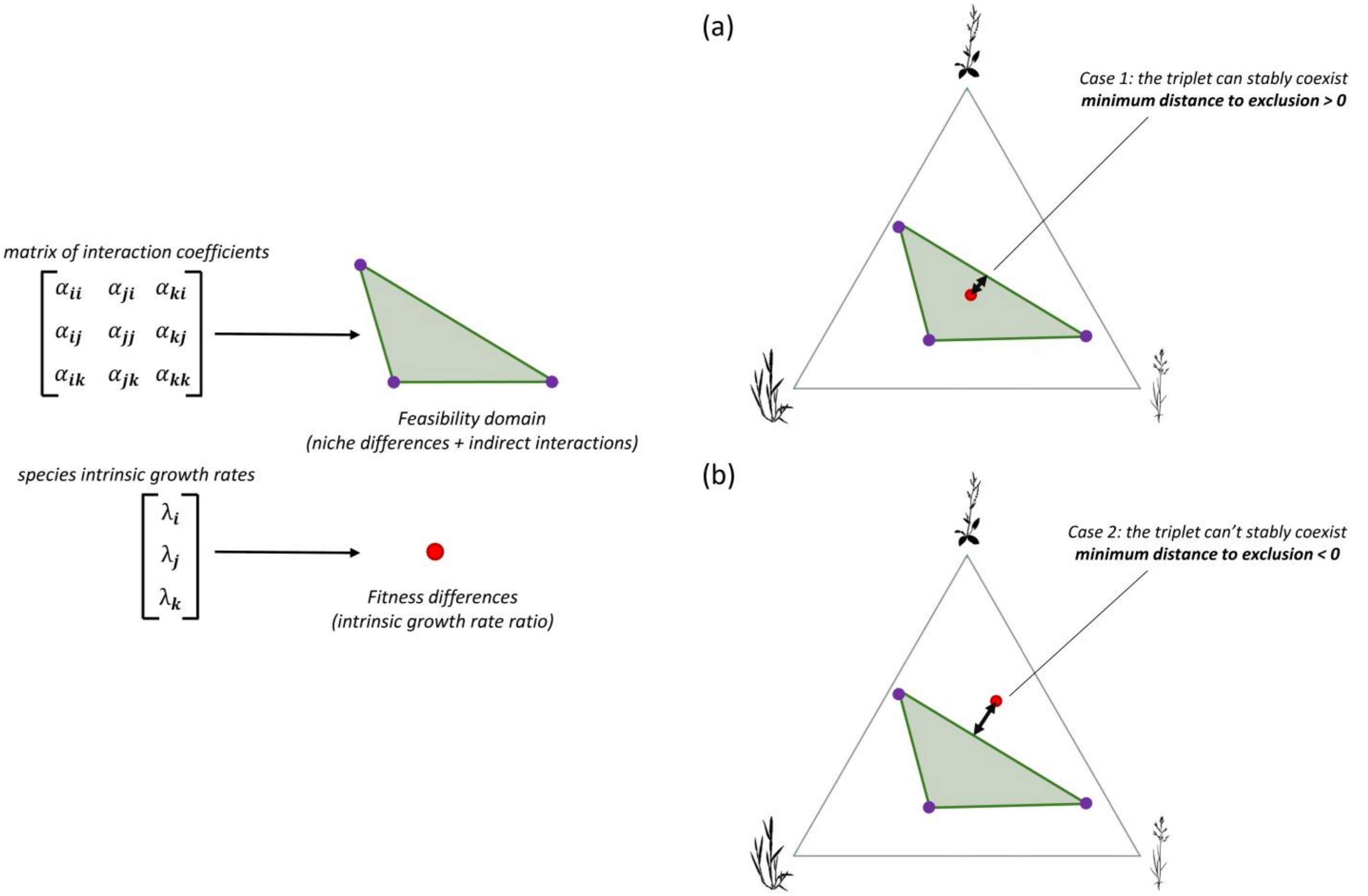


- Figure S10: Structural stability approach, calculations of coexistence mechanisms and of the continuous measure of stable coexistence: the minimum distance to exclusions. This figure represents how the structural stability approach uses the matrix of species interactions and the intrinsic growth rates to project in a geometrical space the feasibility domain (niche differences and indirect interactions) and the fitness differences for a community of three species. The minimum distance to exclusion is then the smallest orthogonal distance of the fitness differences vector (red point) to the closest edge of the pairwise feasibility domain (green triangle). In (a) the vector of fitness differences falls inside the pairwise feasibility domain, the minimum distance to exclusion is positive and the three species are predicted to stably coexist. On the contrary in (b), the vector of fitness differences falls outside of the feasibility domain, the minimum distance to exclusion is negative and the three species are not predicted to stably coexist.

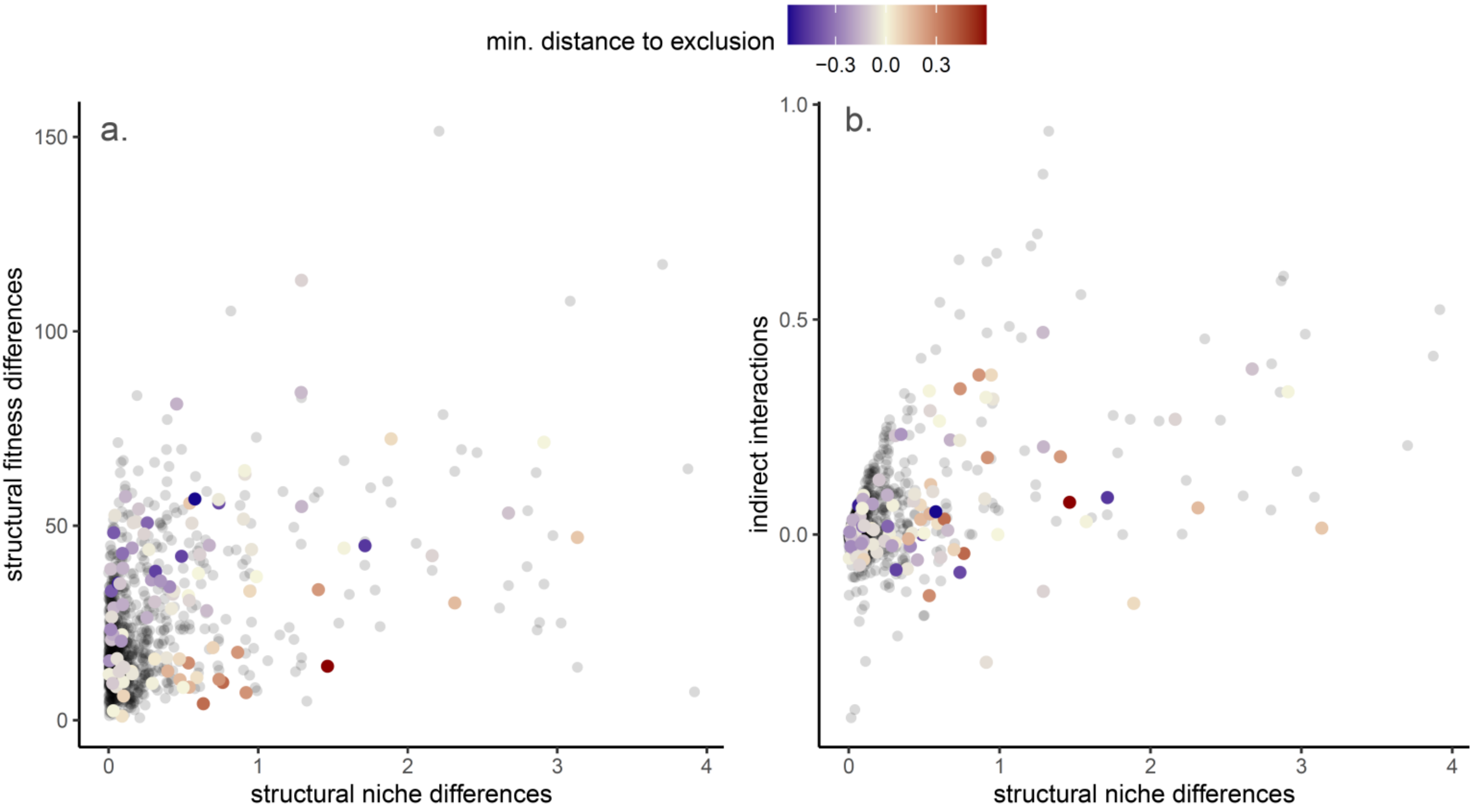


- Figure S11: Distribution of structural niche differences and (a) structural fitness differences and (b) indirect interactions for all possible combination of three out of 18 species sampled in PaNDiv experiment (in black, see Method), and for the 48 combinations of three species selected in the design of this study (n = 12 species) the colours of these 48 points represent their minimum distance to exclusion (blue = non coexisting community, red = coexisting community).

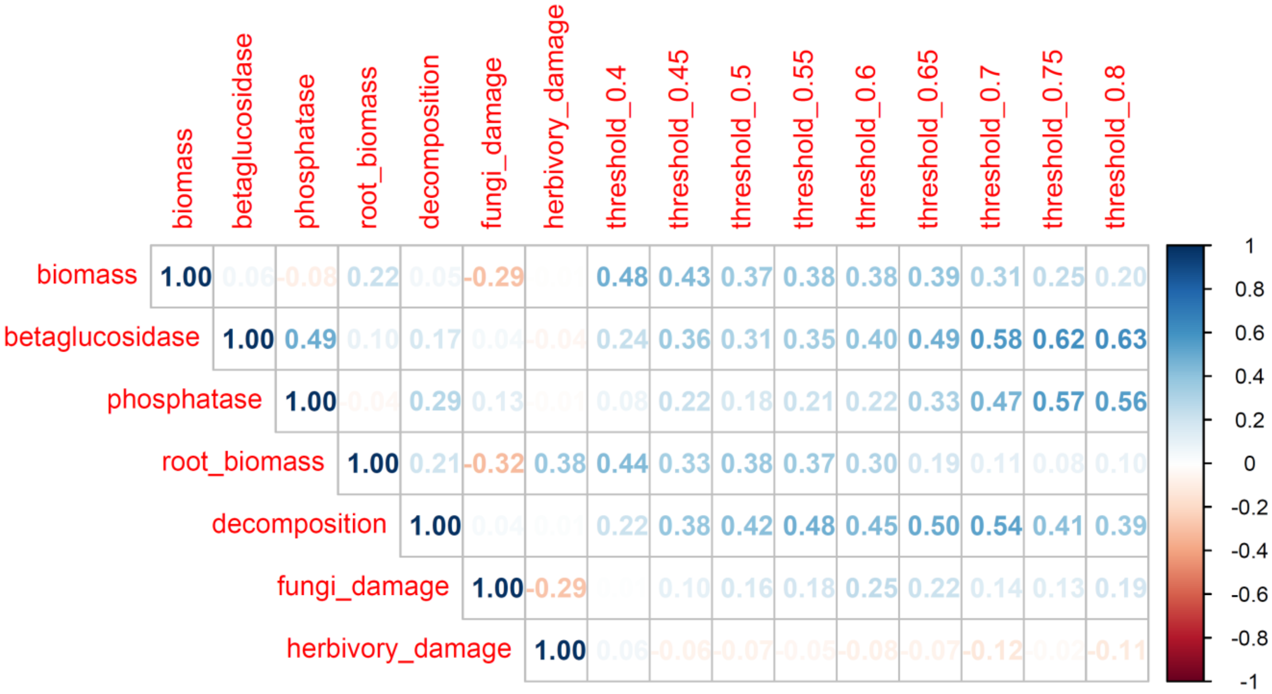


- Figure S12: Correlation plot between each 7 individual functions and the 9 thresholds of multifunctionality.
